# Genomic and Transcriptomic Characterization of Carbohydrate-Active Enzymes in the Anaerobic Fungus *Neocallimastix cameroonii* var. constans

**DOI:** 10.1101/2025.01.29.635548

**Authors:** Elaina M. Blair, Tejas A. Navaratna, Colleen B. Ahern, Ramya Ragunathan, Jennifer L. Brown, Stephen J. Mondo, Anna Lipzen, Radwa A. Hanafy, Kurt LaButti, Jayson Talag, Kerrie Barry, Mansi Chovatia, Mei Wang, Jessy Gonzalez, Xuefeng Peng, Igor V. Grigoriev, Michelle A. O’Malley

**Affiliations:** Department of Chemical Engineering, University of California, Santa Barbara, CA 93106, USA; U.S. Department of Energy Joint Genome Institute, Lawrence Berkeley National Laboratory, Berkeley, CA 94720, USA; Department of Chemical and Biomolecular Engineering, University of Delaware, Newark, DE 19716, USA; Arizona Genomics Institute, School of Plant Sciences, University of Arizona, Tucson, AZ 85721, USA; Department of Plant and Microbial Biology, University of California Berkeley, Berkeley, CA 94720, USA; Joint BioEnergy Institute (JBEI), Emeryville, CA, 94608, USA; Department of Bioengineering, University of California, Santa Barbara, CA 93106, USA

**Keywords:** anaerobic fungi, lignocellulose, RNA-seq, CAZymes, enzyme

## Abstract

Anaerobic gut fungi effectively degrade lignocellulose in the guts of large herbivores, but there remains a limited number of isolated, publicly available, and sequenced strains that impede our understanding of the role of anaerobic fungi within microbial communities. We isolated and characterized a new fungal isolate, *Neocallimastix cameroonii* var. constans, providing a transcriptomic and genomic understanding of its ability to degrade diverse carbohydrates. This anaerobic fungal strain was stably cultivated for multiple years *in vitro* among members of an initial enrichment microbial community derived from goat feces, and it demonstrated the ability to pair with other microbial members, namely archaeal methanogens to produce methane from lignocellulose. Genomic analysis revealed a higher number of predicted carbohydrate-active enzymes encoded in the *N. cameroonii* var. constans genome compared to most other sequenced anaerobic fungi. The carbohydrate-active enzyme profile for this isolate contained 660 glycoside hydrolases, 160 carbohydrate esterases, 194 glycosyltrasferases, and 85 polysaccharide lyases. Differential gene expression analysis showed the upregulation of thousands of genes (including predicted carbohydrate-active enzymes) when *N. cameroonii* var. constans was grown on lignocellulose (reed canary grass) compared to less complex substrates, such as cellulose (filter paper), cellobiose, and glucose. AlphaFold was used to predict functions of transcriptionally active yet poorly annotated genes, revealing feruloyl esterases that likely play an important role in lignocellulose degradation by anaerobic fungi. The combination of this strain’s genomic and transcriptomic characterization, omics-informed structural prediction, and robustness in microbial co-culture make it a well-suited platform to conduct future investigations into bioprocessing and enzyme discovery.

## Introduction

Anaerobic gut fungi are robust lignocellulose degraders, with extensive enzymatic machinery involved in breaking down plant biomass (Hooker et al. 2019). They are pivotal members of large herbivore gut microbiomes where they interact with members of a diverse microbial community, including methanogens that catabolize fungal fermentation products (Li et al. 2021). Several anaerobic gut fungal genera have been cultivated, and the number of isolated strains continues to increase (Hanafy et al. 2022). Still, the genomes of anaerobic gut fungi encode many proteins of yet unknown function (Gruninger et al. 2018; Swift, Louie, et al. 2021), which may bestow these organisms with additional enzymatic mechanisms than those that can be predicted from current annotation tools alone (Lankiewicz et al. 2022).

Complicating the sequencing-based analysis of anaerobic gut fungi is their extremely high adenine-thymine (AT) content and many repeat regions in their genomes (Brownlee 1989; Wilken et al. 2021), which together make it challenging to assemble high-quality genomes (Edwards et al. 2017). However, long-read sequencing has vastly advanced the potential for obtaining high-quality genomes for these microbes (Youssef et al. 2013; Wilken et al. 2019), enabling the annotation of genes and even the construction of genome scale metabolic models (Wilken et al. 2021). Transcriptomic analyses have improved fungal genome annotations and have identified a trove of potential genes involved in lignocellulose breakdown. For example, exploiting catabolite repression patterns across a range of anaerobic fungal strains has revealed enzymes associated with fungal cellulosomes (Solomon et al. 2016; Henske et al. 2017), sugar and carbohydrate transporters (Seppälä et al. 2019), and stress response genes associated with metabolic reprogramming (Swift, Malinov, et al. 2021). This has accelerated efforts to characterize gut fungal enzymes and sugar transporters via heterologous expression (Perli et al. 2021; Podolsky et al. 2021; Liu et al. 2022; Lillington et al. 2023) to link sequence to function. Moreover, while very few anaerobic fungal proteins have high-resolution structures (Dementiev et al. 2023), new approaches in cryoEM (Bai et al. 2015) and protein structure prediction via AlphaFold (Jumper et al. 2021) are accelerating the pace of gene characterization from unconventional microbes. Comparative ‘omics’ studies across multiple anaerobic fungal isolates are necessary to identify groups of genes that are uniquely conserved within this clade, which provide a critical starting point to unmask gene function. However, only 7 strains of anaerobic fungi have published genomes with accompanied transcriptomics characterization (Hanafy et al. 2023), which severely hampers efforts aimed at linking gene sequence to function.

This study reports the isolation and high-quality genome and transcriptome of a new fungal isolate, *Neocallimastix cameroonii* var. constans. This strain was isolated from a laboratory-cultivated anaerobic microbial community derived from goat feces and cultured for multiple years on lignocellulose, with the initial *in vitro* growth and first 10 transfers on alfalfa (Peng et al. 2021). Genomic sequencing, carbohydrate-active enzyme (CAZyme) profiling, phylogenetic analysis, transcriptomic sequencing (RNA-seq), and structure prediction of under-characterized proteins of *N. cameroonii* var. constans were performed on cultures grown on a range of different substrates. These results show that this strain has high CAZyme gene content and is predicted to encode a larger number of glycoside hydrolases compared to most other anaerobic gut fungi sequenced to date. Based on structural predictions, there are additional predicted CAZymes (including feruloyl esterases) encoded in this fungal isolate genome that are missed with more traditional annotation methods. Several feruloyl esterases are predicted in proteins containing dockerin domains, thus supporting the importance of cellulosomes in anaerobic fungal carbohydrate degradation. It is also shown that co-cultures containing *N. cameroonii* var. constans and methanogenic archaea can produce methane, similar to other reported strains of AGF (Cheng et al. 2009; Swift et al. 2019; Leggieri et al. 2021).

## Materials and Methods

### Microbial Enrichment and Roll Tube Isolation of Anaerobic Fungi

The anaerobic fungus *N. cameroonii* var. constans was isolated from a consortium of fungi, methanogens, and bacteria. The consortium was enriched from the feces of a San Clemente Island goat at the Santa Barbara Zoo through extended cultivation on an alfalfa substrate and regular antibiotic treatment with penicillin and streptomycin as described previously (Peng et al. 2021). Shortly after the 10-passage consecutive batch culture enrichment (Peng et al. 2021), the substrate was changed to reed canary grass, and the community was cultivated for approximately 3 years via regular anaerobic passage in Hungate tubes (9 mL MC– media (Peng et al. 2018) with approximately 0.1 g dried, milled reed canary grass) with a transfer every 3-4 days following inoculation and growth monitoring via headspace pressure monitoring (Haitjema et al. 2014). Cultures were passaged using a sterile syringe-needle technique, with 1 mL of growing culture transferred to fresh media each passage.

Roll tube isolation was used to select for a single anaerobic fungal isolate (Haitjema et al. 2014). After 3 years of cultivation, 1 mL of the consortium culture was inoculated into 9 mL Medium C (Davies et al. 1993) in a Hungate tube supplemented with 0.1 mL chloramphenicol (prepared in 40 vol% molecular grade ethanol; 100 µg/mL final concentration) under anaerobic conditions with dried and milled reed canary grass (approximately 0.1 g) supplied as a carbon source. The consortium was cultivated in this way with inoculation into a fresh Hungate tube every 3-4 days of growth until methane could no longer be detected via gas chromatograph (TRACE 1300, Thermo Fisher Scientific) and the culture did not appear turbid. A fungal isolate was then selected with the roll tube isolation method (Haitjema et al. 2014), with thallus picking performed inside an anaerobic chamber (cat. no. AS-580, Anaerobe Systems, Morgan Hill, California, USA). After subsequent growth observed by pressure production and grass clumping, 0.1 mL of the fungal culture supernatant was used to inoculate an anaerobic roll tube. The roll tube was allowed to grow for approximately 4 days until visible thalli formed on the surface of the agar. A single thallus from a clonal fungus was picked from the agar wall of the roll tube in an anaerobic chamber (cat. no. AS-580, Anaerobe Systems). The thallus was inoculated into a Hungate tube containing 10 mL Medium C with 0.1 mL chloramphenicol (100 µg/mL final concentration) and reed canary grass supplied as a carbon source. The roll tube isolation process was repeated four times to ensure axenic cultivation, alternating between single thallus picking in roll tubes and inoculation into liquid cultures in the presence of chloramphenicol.

### Routine Cultivation and Culture Media

Monocultures of *N. cameroonii* var. constans were routinely grown anaerobically at 39°C in Hungate tubes containing 9 mL of either MC– media (Peng et al. 2018) with vitamin supplement (0.1% v:v, ATCC cat. no. MD-VS, made in house or ordered from ATCC) or Medium C (Davies et al. 1993) and approximately 0.1 g lignocellulosic substrate (either dried, milled reed canary grass or sorghum). Cultures were passaged every 3-5 days, and 1 mL growing culture was used to inoculate fresh media.

### Morphological and Phylogenetic Characterization of the Fungal Isolate

Approximately 10-20 µL of growing fungal culture were prepared on a microscope slide and imaged using a Zeiss Primovert transmitted light microscope for morphological characterization (cat. no. 415510–1101-000, Carl Zeiss Microscopy, Oberkochen, Germany) with a 20x objective and a SPOT Idea 28.2 5-MP camera (SPOT Imaging, Sterling Heights, Michigan, USA). Spot 5.1 software was used to capture images and add scalebars. Fungal zoospores were identified based on cell size (cell bodies ranging from 5-10 μm), circular morphology, and movement with the presence of flagella.

The full ITS1-D1/D2 LSU genomic region was amplified from genomic DNA extracted with the DNeasy PowerSoil Pro kit (Qiagen) according to manufacturer protocol, using primers ITS5 (5’- GGAAGTAAAAGTCGTAACAAGG-3’) and GG-NL4 (5’-TCAACATCCTAAGCGTAGGTA-3’) (Hanafy et al. 2020) and cloned into chemically competent DH5α *E. coli* using the Invitrogen TOPO TA Cloning Kit for Subcloning. The D1/D2 LSU region was amplified from 7 colonies by colony PCR using Phusion polymerase and primers NL1 (5’- GCATATCAATAAGCGGAGGAAAAG-3’) and GG-NL4, and the amplicons were submitted for Sanger sequencing. Sequences were aligned with a reference dataset of anaerobic fungal D1- D2 LSU sequences, obtained from the NCBI-GenBank nr database (Supplemental Table 1), using MUSCLE (Edgar 2004) with default parameters and manually refined in Geneious software. The generated alignment was used for constructing a Maximum Likelihood phylogenetic tree using IQ-TREE 2 (Hoang et al. 2018; Minh et al. 2020), with *Chytriomyces* sp. WB235A as the outgroup. A best fit substitution model (TN+F+G4) was chosen according to the Bayesian Information Criterion. Bootstrap values were calculated based on 1000 replicates. The final tree was visualized and edited using the Interactive Tree of Life (iTOL) platform (Letunic and Bork 2021). Based on combined morphological and ITS analysis, the fungal isolate was named *N. cameroonii* var. constans because of phylogenetic clustering within the *N. cameroonii* clade (Supplemental Figure 1).

### Harvesting Tissue for DNA extractions

Fungal monocultures were grown for three days prior to harvest. They were cultivated in anaerobic serum bottles containing 80 mL MC– media (Peng et al. 2018), with 0.2 µm filtered vitamin supplement (0.1% v:v) and glucose (5 g/L) added post autoclaving. Ten bottles were combined and vacuum filtered through sterile miracloth and rinsed with Millipore water. The fungal mat was then removed from the miracloth with tweezers and put in a 50 mL falcon tube. It was flash frozen in liquid nitrogen and then stored in a -80°C freezer until shipment to the Arizona Genome Institute on dry ice.

### Genomic DNA Extraction and Sequencing

DNA was extracted at the Arizona Genomics Institute using a modified cetyl trimethylammonium bromide (CTAB) protocol. DNA was then sequenced at the Joint Genome Institute (JGI) with the PacBio SEQUEL IIe using the protocol for 6-10kb with BluePippin size selection with 1x1800 minute sequencing movie times. The CCS reads were filtered for artifacts and then assembled with Flye version 2.9-b1768 [-t 32 --pacbio-hifi] (https://github.com/fenderglass/Flye) (Kolmogorov et al. 2019) and subsequently polished with two rounds of RACON version 1.4.13 racon [-u -t 36] (https://github.com/lbcb-sci/racon) (Vaser et al. 2017). The final genome assembly was annotated using the JGI Annotation pipeline (Grigoriev et al. 2014). GC% was quantified using the infoseq command in EMBOSS version 6.6.0 (Rice et al. 2000) for unmasked assemblies and CDS-only FASTA files downloaded from MycoCosm (Grigoriev et al. 2014).

### CAZyme Gene Annotation

Genomes were accessed and downloaded from MycoCosm (Grigoriev et al. 2014). For comparisons of CAZyme genome content and predictions across the fungal kingdom, dbCAN-sub (Zheng et al. 2023) was run using default parameters on all genomes on the California Nanosystems Institute’s Pod cluster at UCSB, and the outputs containing CAZyme annotations and substrate predictions were parsed using custom R scripts. All unique substrates were summed during tabulation, even when multiple substrates were predicted for a given CAZyme. The prediction of multiple substrates may represent enzyme promiscuity, or alternately, model ambiguity requiring experimental determination.

For tabulation of dockerin and scaffoldin domains in *N. camerooni* var. constans, which is outside the scope of dbCAN-sub, HMMER searches (hmmer.org, version 3.1b2) (Finn et al. 2011) of the CDS aa.fasta file downloaded from MycoCosm were carried out using PF02013.hmm for dockerin and cohesin3.hmm. Annotations for CAZymes, for the enumeration of dockerins and carbohydrate-binding modules fused to enzymatic domains in the genome of *N. camerooni var. constans* were downloaded from MycoCosm (Grigoriev et al. 2014) and parsed using custom R scripts. All HMMER search models, CAZyme list, and open reading frame file are available on https://github.com/O-Malley-Lab/N_var_constans.

### Cultivation for RNA Extraction, Sequencing, and Assembly

Hungate tubes containing 9 mL MC– media (Peng et al. 2018), vitamin supplement (0.1% v:v, made in house from ATCC cat. no. MD-VS recipe), and either glucose (5 g/L), cellobiose (5 g/L), filter paper (0.1 g, cut in strips), or reed canary grass (0.1 g, 1 mm sieved) (4 biological replicates of each condition) were inoculated with 1 mL from a 3-day old fungal culture grown on MC– with reed canary grass. Cultures were grown for 2-3 days and were then poured into 50 mL falcon tubes containing 10 mL RNAlater, made in-house (Malmstrom 2015; Erster et al. 2021) and centrifuged for 30 minutes at 10,000 x g and 4°C using a fixed angle rotor (Eppendorf 5810 R, rotor F-34-6-38). After centrifugation, the supernatant was decanted. The pellets were flash frozen in liquid nitrogen and then stored in a -80°C freezer until RNA extraction.

RNA extractions were performed using the Qiagen RNeasy Mini Kit, following the manufacturer-provided protocol: “Purification of total RNA from plant cells and tissues and filamentous fungi” with the addition of the optional DNase on-column digest. Cells were lysed via liquid nitrogen grinding, and a QIAshredder was used to homogenize each sample. RNA was eluted in 50 µL of RNAse-free water, and the eluent was then re-eluted through the column to concentrate the RNA. RNA concentration was checked on the Invitrogen Qubit 2.0 fluorometer, with all samples >25 ng/µL. Quality was evaluated with the Agilent 2200 TapeStation or 2100 Bioanalyzer – all samples had a RIN or RINe score above 7.

Extracted RNA was sent to the JGI and sequenced using the Illumina NovaSeq S4 with run type 2x151 bp. PCR was employed to make stranded sequencing libraries. The JGI used their pipeline for quality control; BBDuk was used to remove artifacts from the 3’ end (settings: kmer=25, 1 mismatch allowed, phred trimming at Q6) and reads with Ns, PhiX, or RNA spike-in. Short reads were also removed (<1/3 of initial read length and/or <25 bp). Alignment to the newly sequenced genome was performed with HISAT2 (v. 2.2.1) (Kim et al. 2015); output files were sorted and indexed using SAMtools v. 1.7 (Li et al. 2009). deepTools (v. 3.1) was used to calculate strand-specific coverage (Ramírez et al. 2014), and raw gene counts were calculated with featureCounts v. 1.5.2 (settings: -s 2 -p --primary options) (Liao et al. 2014).

### Differential Gene Expression Analysis, Multidimensional Scaling (MDS) Plot and Heatmap

The raw gene count matrix generated by featureCounts (see above) was used as the input for DESeq2 v. 1.40.2 (Love et al. 2014). Genes with zero counts were removed prior to running DESeq2. After using the DESeq command, pairwise conditions were compared using the contrast function. Plots showing significantly regulated genes between the pairwise comparisons were made using the DESeq2 plotMA command with alpha set to 0.05 (Supplemental Figure 2). Except for in this supplemental figure, genes with an average transcripts per million (TPM) value less than two for at least one condition were excluded from further analysis. Significance in analyses was set to a p-adjusted value <0.05.

The multidimensional scaling (MDS) plot was created using the edgeR package in R (Robinson et al. 2010). After applying the TPM cutoff as in the preceding paragraph, the raw gene count matrix was normalized using trimmed-mean-of-M-values (TMM) normalization. The MDS plot was then generated from these counts using Euclidean distance via the plotMDS function.

The pheatmap package in R was used to generate the differential expression heatmaps. P-values from the filtered dataset were used to generate a heatmap of 100 genes with the minimum of p-values across the three comparative conditions (cellobiose vs. glucose, filter paper vs. glucose, and reed canary grass vs. glucose). The p-values for the condition of reed canary grass vs. glucose dominate the minimum p-values used in this analysis. Additional heatmaps were therefore created using the minimum p-values for cellobiose vs. glucose only (Supplemental Figure 3), filter paper vs. glucose only (Supplemental Figure 4), and reed canary grass vs. glucose only (Supplemental Figure 5). Gene-level annotations for CAZymes were downloaded from MycoCosm (Grigoriev et al. 2014).

### CAZyme Structure Predictions

Structures of highly differentially regulated proteins containing dockerin and/or CBM domains but lacking further enzymatic annotations were predicted using AlphaFold (Jumper et al. 2021), which was implemented on California Nanosystems Institute’s high performance computing clusters. Proteins of interest were selected by sorting predicted CAZyme genes that had only dockerin and/or CBM domain annotations (MycoCosm) by lowest adjusted p-values for differential gene expression. The highest ranked structure prediction from AlphaFold for each protein was used for further analyses. The per-residue local distance difference test (pLDDT) confidence scores for the protein structure models were retrieved from the B-factor field of the coordinate section of the output pdb file and the final structures were visualized using UCSF ChimeraX (Meng et al. 2023). Structural homology searches for annotation were carried out using the Foldseek web interface (van Kempen et al. 2023). The protein sequences were also run on InterPro (Blum et al. 2024) for comparison.

### Roll Tube Isolation and Characterization of Methanogen from Enrichment Consortium

A *Methanobrevibacter* sp. methanogen was isolated after 3 years of cultivation from the same enrichment consortium as *N. cameroonii* var. constans. Methanogen media was used to prepare roll tubes – this culture medium is MC– medium (Peng et al. 2018) with the following additions per liter: 10 mL of 100x trace elements solution (Lowe et al. 1985), 2 g sodium acetate anhydrous, and 4 g sodium formate. Methanogen media was bubbled with carbon dioxide prior to aliquoting, and media was aliquoted under a flow of 80% hydrogen 20% carbon dioxide. Roll tubes were prepared with 5 mL methanogen media and 0.1 g agar. 0.1 mL of vitamin supplement was added to the roll tubes post autoclaving. For inoculation, 0.1 mL of the enrichment culture was added via sterile syringe and needle into the roll tube. Roll tubes were incubated at 39°C for 7–10 days to allow for methanogen colony formation. In an anaerobic chamber (cat. no. AS-580, Anaerobe Systems), roll tubes were opened, and a colony was selected and used to inoculate a tube of liquid methanogen media. This tube was incubated at 39°C for 7–10 days, after which it was used to inoculate a new roll tube. The roll tube isolation process was repeated 2 more times (for a total of 3 roll tube isolations). Each liquid culture was evaluated for and showed methane production.

For characterization of this isolate, the 16S rRNA gene was amplified from gDNA of two liquid cultures and sequenced using Sanger sequencing. The primers used were: Met86F (5′- GCTCAGTAACACGTGG-3′) and Met1340R (5′-CGGTGTGTGCAAGGAG-3′) (Wright and Pimm 2003). When these sequences were run on the NCBI BLAST webserver (blastn, default parameters) (Camacho et al. 2009), top hits belonged to *Methanobrevibacter* spp., and sequence similarity was >99% identity in many cases; hence, this isolate is referred to as *Methanobrevibacter* sp.

### Co-cultivation of Anaerobic Fungi with Methanogenic Archaea and Methane Analysis

*Methanobrevibacter smithii* (DSM 861), *Methanobrevibacter thaueri* (DSM 11995), and *Methanobacterium bryantii* (DSM 863) were obtained from the DSMZ culture collection. These methanogens and the methanogen isolated in this study were revived from cryostocks and cultivated at 39°C on methanogen media described above, except trace elements solution followed the ATCC MD-TMS recipe (made in-house). Vitamin supplement (0.1% v:v) was added post autoclaving to a Hungate tube containing 0.1 g dry, milled reed canary grass and 9 mL Medium C (Davies et al. 1993), 0.5 mL of methanogen-containing culture (see above) and 0.5 mL of anaerobic fungi-containing Medium C culture were added. Cultures were incubated at 39°C; after 4 days of co-culture growth, gas pressures were measured via pressure transducer, and 100 μL of headspace gas was injected into a Shimadzu GC-14A equipped with a flame ionization detector for methane quantification by comparison with a standard curve.

## Results and Discussion

### *Neocallimastix cameroonii* var. constans was Isolated from a Mixed Goat Fecal Consortium

*N. cameroonii* var. constans was isolated as part of an extensive enrichment study, where communities from goat feces were cultivated on different biomass substrates and characterized via whole-genome shotgun sequencing (Peng et al. 2021). *Neocallimastix* was the most abundant anaerobic fungal genus cultivated on the three different lignocellulosic substrates tested, and out of the 18 fungal metagenome-assembled genomes produced from the Peng *et al*. study, 12 are likely the same species as *Neocallimastix californiae* (Peng et al. 2021) (also known as *Neocallimastix camerooni* var. californiae). The fungal isolate, *N. cameroonii* var. constans, is of particular interest because it belongs to that species and contributed to those MAGs, and because prior to isolation, it was stably cultivated for 3 years in a consortium composed mainly of fungi and methanogens (Peng et al. 2021). Given its longevity in cultivation compared to other anaerobic fungi, its capacity to survive cryopreservation, and its ability to stably pair with other anaerobes, it represents a robust strain amenable to long-term culture that was characterized further in this study.

This isolate, *N. cameroonii* var. constans (characterized as a variant of the *Cameroonii/Californiae* species), is shown in micrograph in Figure 1, and its fungal structure, including sporangia, rhizoids, and zoospores, are all consistent with assignment to the *Neocallimastix* genus. *Neocallimastix* fungi have an extensive rhizoidal network and monocentric thalli (Orpin 1975; Hess et al. 2020). Like other anaerobic fungi, *N. cameroonii* var. constans has a genome of 187 Mbp that is very AT-rich. There is a broad range of copy number variation among genes, but no evidence of specific polyploidy. The GC % of the entire genome is 18.2%, which is within the 16-22% range reported for another anaerobic fungi (Wilken et al. 2020). For coding regions, the GC content is 28.1%. Sequencing statistics are presented in Table 1. *N. cameroonii* var. constans as isolated here was likely to be a member of the species complex containing other conspecific *Neocallimastix* variants in the enriched microbial goat communities and shares >99.5% ITS2 sequence identity with several *Neocallimastix* MAGs from Peng *et al* (Peng et al. 2021). To better understand the relationship between these species in relation to other *Neocallimastigomycota,* we analyzed large ribosomal subunit (LSU) sequences (Hanafy et al. 2020) and built a phylogenic tree (Supplemental Figure 1).

**Figure 1.**
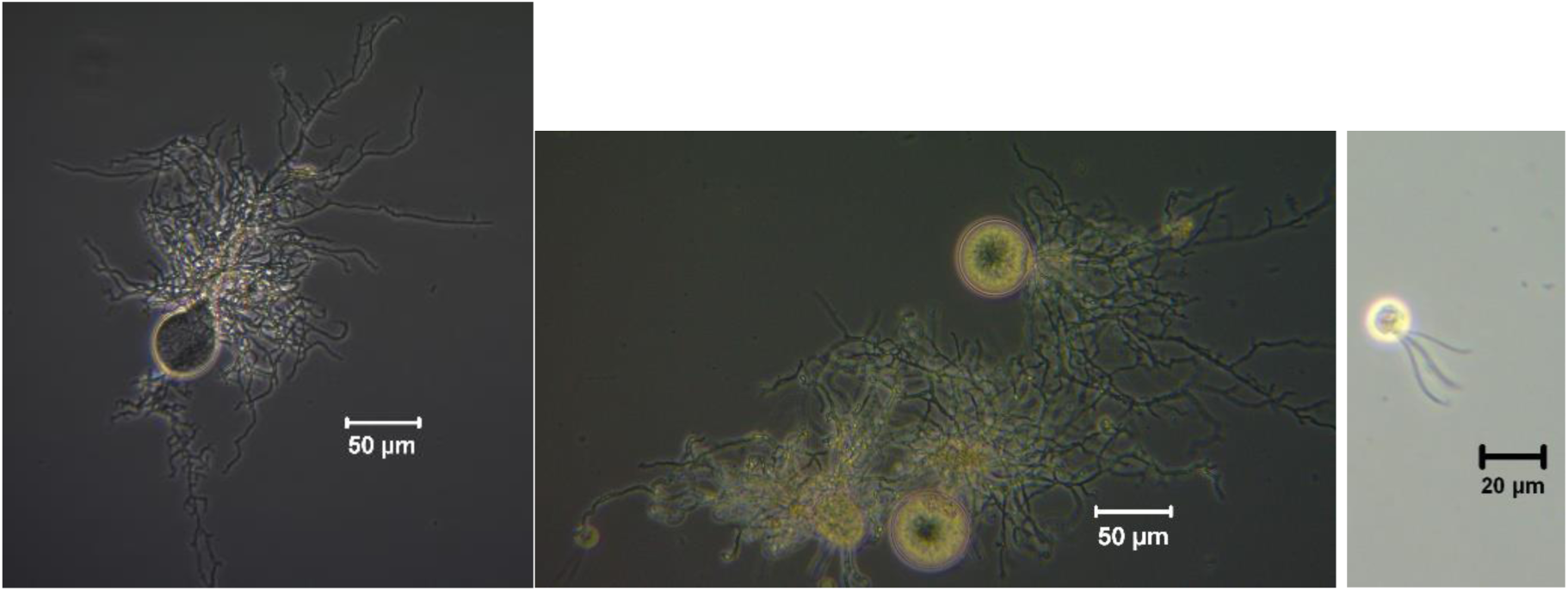
Microscopy shows monocentric thalli (A,B), rhizoids (A,B), and polyflagellate zoospore (C) morphology of *N. cameroonii* var. constans.

**Table 1.**
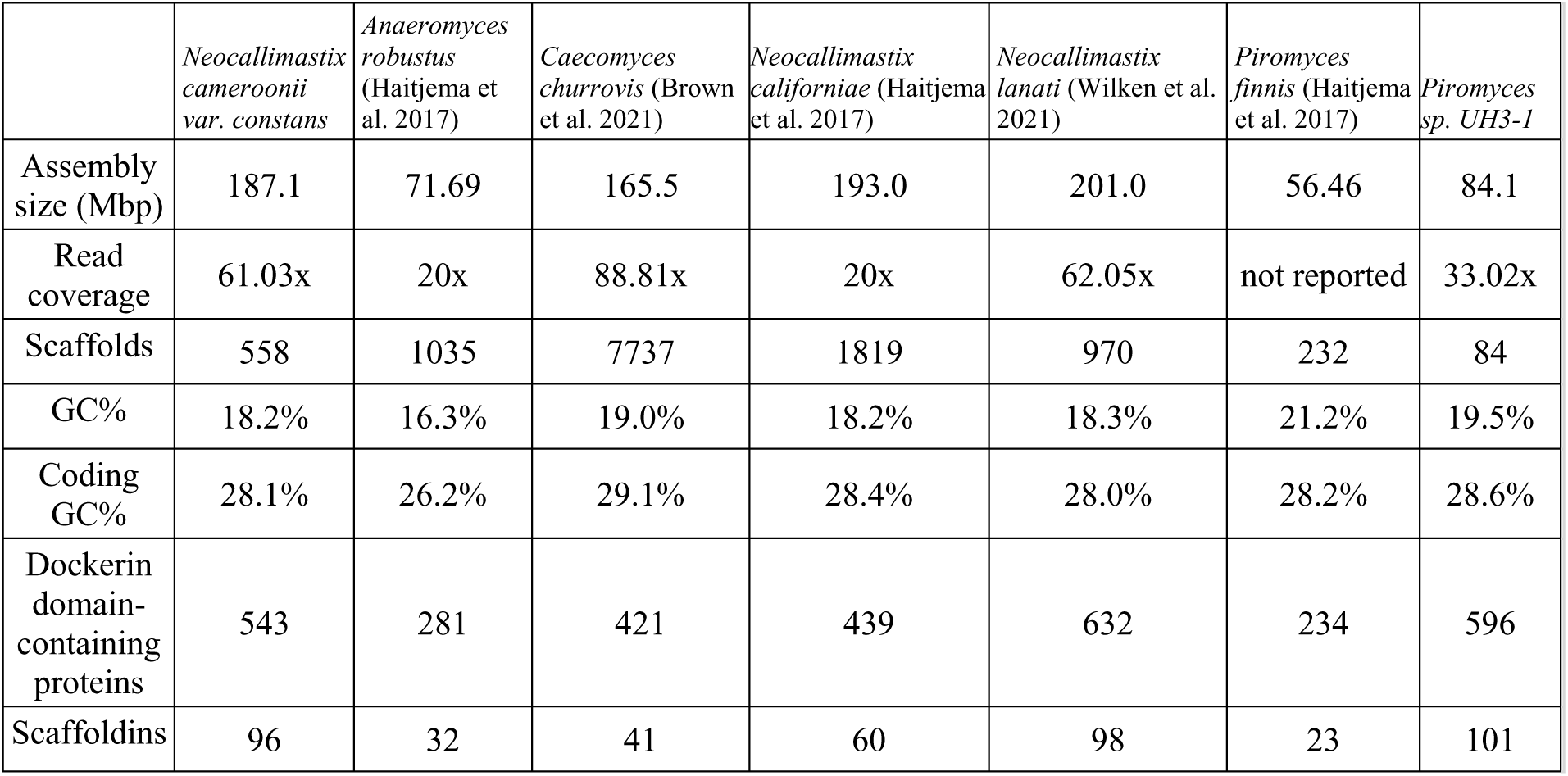
Genome statistics for *N. cameroonii* var. constans and select other anaerobic fungi.

### Numerous Predicted CAZymes are Encoded in the *N. cameroonii* var. constans Genome

Characterizing the CAZyme repertoire for a given fungus is a valuable tool for predicting its ecology. *B. dendrobatidis,* the well-publicized frog pathogen, is notable for its genomic expansion of chitin-binding domains (Abramyan and Stajich 2012). *M. rebaudengoi,* which grows on dead and decaying plant matter in forests, contains 118 putative lignin-active enzymes. Similarly, anaerobic fungal isolates have many enzymes targeted to a broad range of carbohydrates, including xylan, beta-glucan, chitin, beta-mannan, pectin, beta-galactan, and starch.

Using dbCAN3 (Zheng et al. 2023), we annotated CAZyme domains for comparison across fungi from various phyla. The closely related strain *N. californiae* (Haitjema et al. 2017) had very similar CAZyme composition and count as well as genome size (Figure 2A). While the mushroom-forming basidiomycete *Mycena rebaudengoi* (Grigoriev et al. 2014) contained more total CAZymes, a large fraction of these were categorized as auxiliary activities, which consist of redox enzymes including lignin-active lytic polysaccharide mono-oxygenases among others (Cantarel et al. 2009). These observations confirm that anaerobic fungi contain the largest genomic repertoire of core CAZymes, consisting of carbohydrate esterases, glycoside hydrolases, glycosyltransferases, and polysaccharide lyases (Figure 2).

**Figure 2.** (A) *N. cameroonii* var. constans has among the highest CAZyme domain content compared to other sequenced fungi. Abbreviations are as follows: *A. robustus* (*Anaeromyces robustus*), *A. flavus* (*Aspergillus flavus*), *A. oryzae* (*Aspergillus oryzae*), *B. dendrobatidis* (*Batrachochytrium dendrobatidis*), *C. churrovis* (*Caecomyces churrovis*), *C. albicans* (*Candida albicans*), *G. prolifera* (*Gonapodya prolifera*), *M. rebaudengoi* (*Mycena rebaudengoi*), *N. var. constans* (*Neocallimastix cameroonii* var. constans), *N. californiae* (*Neocallimastix californiae*), *P. sp C1A* (*Pecaromyces sp. C1A*), *P. finnis* (*Piromyces finnis*), *R. pusillus* (*Rhizomucor pusillus*), *S. cerevisiae* (*Saccharomyces cerevisiae*), *S. pombe* (*Schizosaccharomyces pombe*), *S. punctatus* (*Spizellomyces punctatus*), *T. reesei* (*Trichoderma reesei*). Proteins were annotated using dbCAN3. (B) Substrate prediction for CAZymes identified in *N. cameroonii* var. constans using dbCAN-sub. For clarity, the 30% of enzymes lacking a substrate prediction were omitted for the chart.

With dbCAN-sub (Zheng et al. 2023), we further enumerated putative substrates for identified CAZymes in *N. cameroonii* var. constans, as well as in other fungi for comparative analysis. Substrate analysis confirmed extensive lignocellulose-degrading activity, with a near-majority (Figure 2) of CAZymes targeted at xylan, and beta-glucan and cellulose being two other highly represented substrates.

### CAZymes Contain a Broad Range of Cellulosome-Associated Features

Anaerobic fungi produce cellulosomes, which are multi-enzyme free or surface-anchored assemblies more widely studied in many cellulolytic bacteria (Doi et al. 2003; Artzi et al. 2017). Across species of anaerobic fungi, conserved dockerin domains are found in many CAZymes (Nagy et al. 2007), and these have been shown to play important roles in cellulosome assembly and biomass breakdown (Haitjema et al. 2017; Gilmore et al. 2020). In *N. cameroonii* var. constans, dockerin and carbohydrate-binding module (CBM) motifs are widespread, with the annotated genome containing 543 dockerin-containing proteins and 96 scaffoldins (Table 1). The distribution of dockerin domains also shows a marked CAZyme-class dependence (Table 2). Consistent with their presumed roles in the synthesis, as opposed to catabolism of carbohydrates, glycosyltransferases (GTs) in *N. cameroonii* var. constans are bereft of dockerins or CBMs (Table 2). Glycosyl hydrolases (GHs), however, frequently contain multiple dockerin domains (Supplemental Figure 6), indicating key roles in lignocellulosic breakdown.

**Table 2.** Cellulosome and CAZyme annotations in *N. camerooni var. constans.* Annotations downloaded from MycoCosm were used to quantify the fraction of CAZyme-containing open reading frames that also contained dockerins and/or carbohydrate binding domains (CBMs). Note that Table 2 represents protein counts, not domain counts; ie. some proteins have multiple domains. Statistics for six other anaerobic fungi are presented in Supplemental Table 17.

| CAZyme class | Protein count | Proteins containing dockerin domains | Fraction containing dockerin domains | Proteins with annotated CBM | Fraction with CBM |
| --- | --- | --- | --- | --- | --- |
| Carbohydrate esterase (CE) | 160 | 20 | 0.13 | 64 | 0.40 |
| Glycoside hydrolase (GH) | 660 | 208 | 0.32 | 155 | 0.23 |
| Glycosyltransferase (GT) | 194 | 0 | 0 | 0 | 0.00 |
| Polysaccharide lyase (PL) | 85 | 2 | 0.02 | 34 | 0.40 |

Since numerous genes found in anaerobic gut fungal genomes encode proteins of unknown function (Lankiewicz et al. 2022), such as those annotated as “hypothetical proteins,” it was hypothesized that there are unannotated CAZymes in the *N. cameroonii* var. constans genome. Specifically, many predicted CAZyme genes are annotated as containing one or more dockerin domains, but the majority of the protein length lacks specific enzymatic annotations. AlphaFold (Jumper et al. 2021) was used to better annotate proteins that were highly differentially regulated during growth on lignocellulosic substrate. Multiple proteins were identified that appear to contain a feruloyl esterase domain along with dockerin domain(s), with some also containing a carbohydrate-binding module 13 (CBM13) (Figure 3). The CBM13 domain was first identified in plant lectins and can bind to sugars multivalently (Fujimoto 2013). In these proteins from *N. cameroonii* var. constans, sequence-based annotation only revealed the possible existence of the superfamily alpha/beta hydrolase fold (Figure 3), which encompasses a wide range of catalytic and non-catalytic proteins across multiple domains of life (Holmquist 2000).

**Figure 3.**
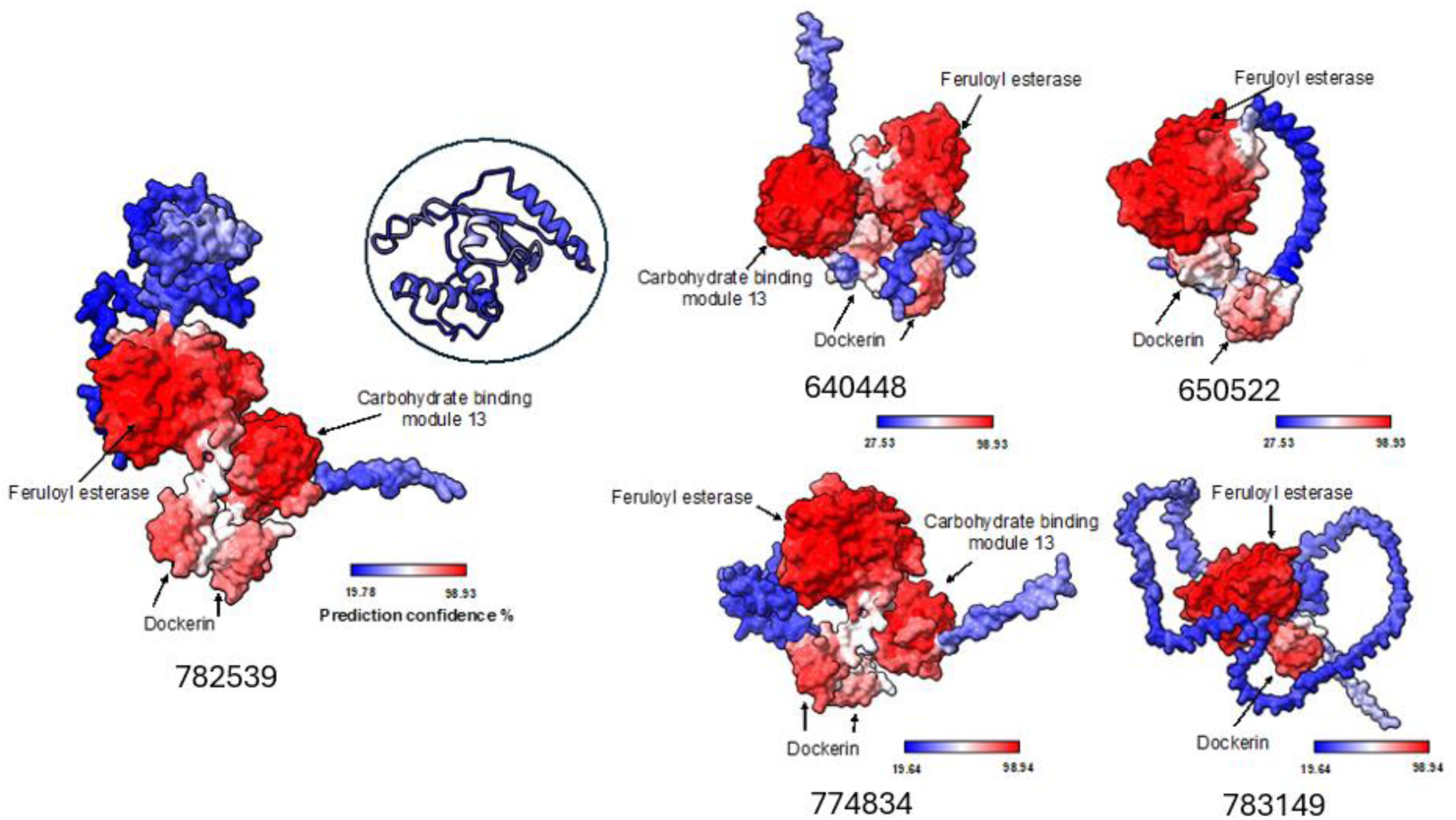
AlphaFold predicted structures show additional unannotated enzyme domains encoded by genes that were highly upregulated when cultures of *N. cameroonii* var. constans were grown on reed canary grass. Predicted structures for MycoCosm ProteinIDs: 782539, 640448, 650552, 774834, and 783149 are shown. Structural homology searching with Foldseek predicts that these proteins are feruloyl esterases. Supplemental Table 2 lists adjusted p-values and log fold change values for pairwise comparisons of these genes when cultures were grown on different substrates.

AlphaFold (Jumper et al. 2021) and FoldSeek (van Kempen et al. 2023) predicted the existence of feruloyl esterase domains in several under-annotated genes in this genome (Figure 3). Feruloyl esterases catalyze bond scission between lignin and polysaccharides, an essential step in the metabolism of plant matter by anaerobic fungi (Dilokpimol et al. 2016). Furthermore, some level of structural (and sequence) diversity exists among these feruloyl esterases, with proteins containing a variable number of dockerins, carbohydrate-binding modules, and most interestingly, a lid-like domain (Figure 3) which has been implicated in substrate binding (Suzuki et al. 2014; Perez-Garcia et al. 2023). However, this putative lid-like domain was not predicted with much confidence by AlphaFold (pLDDT<50), though it seems to have some structural elements (4 alpha helices and 1 beta sheet), indicating it might be only structured in a multiprotein complex or stable under certain conditions such as in presence of calcium or substrate, which has been reported previously (Suzuki et al. 2014). A different low confidence element is predicted in protein 774834 (Figure 3 and Supplemental Table 2). Additionally, these predicted feruloyl esterases are upregulated in presence of lignocellulosic biomass, but not all are upregulated in presence of cellulose-only filter paper (Supplemental Table 2), increasing confidence that these enzymes may be involved in the breaking of bonds at the interface of lignin and polysaccharides. The FoldSeek prediction of feruloyl esterase domains in many of these is consistent with HMM-based annotation by PANTHER (Thomas et al. 2022) even though Superfamily searches (Pandurangan et al. 2019) only returned broad alpha-beta hydrolase classifications. The structure-based matching reported here shows broad core domain conservation but also reveals diverse additional protein modules that may be involved in modulating activity and/or localization. The sequence-level diversity of annotated and unannotated feruloyl esterases in this anerobic fungus genome is consistent with recent experimental support for multi-domain feruloyl esterase-containing proteins in other anaerobic fungi (Shi et al. 2025). While the results presented here are primarily predictions of structure, they invite additional experimental efforts to further identify and characterize these putative feruloyl esterases with regards to protein abundance and substrate specificity.

### Substrate Complexity Determines *N. cameroonii* var. constans CAZyme Expression

Multidimensional scaling (MDS) analysis showed that transcriptomic data from samples of *N. cameroonii* var. constans clustered significantly based on the substrate on which they were cultured (Figures 3–4). The *N. cameroonii* var. constans samples grown on glucose, filter paper, and reed canary grass clustered far from each other on the MDS plot, while the samples grown on cellobiose clustered closely with those grown on glucose. As cellobiose is a disaccharide of glucose, *N. cameroonii* var. constans experiences similar gene expression in response to glucose and cellobiose.

**Figure 4.**
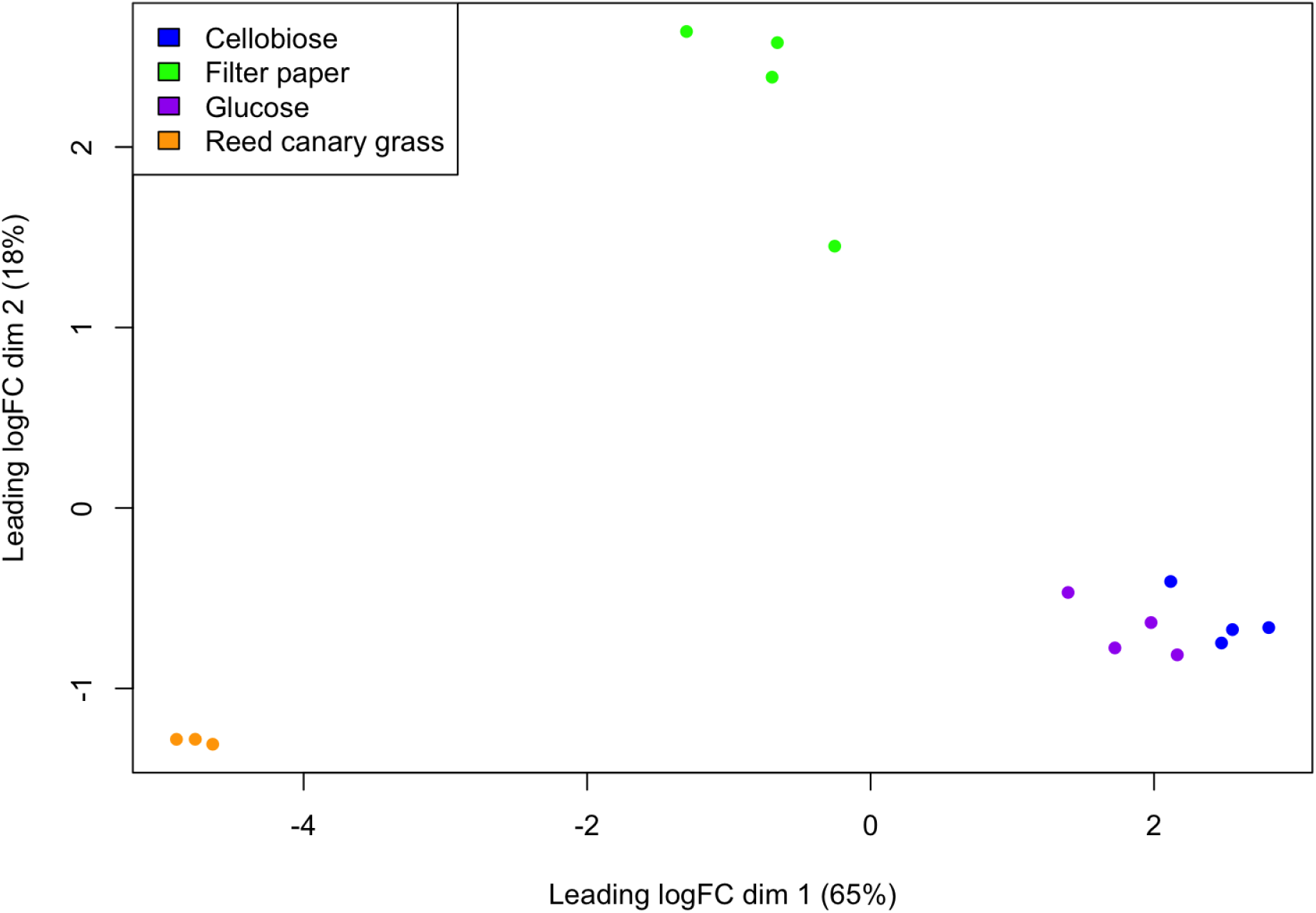
A multidimensional scaling (MDS) plot of *N. cameroonii* var. constans growth on 4 substrates shows that gene expression highly clusters by substrate type. The leading log-fold-change (leading logFC) is the root-mean-square average of the top 500 largest log2-fold-changes between each sample pair.

As expected, we found that CAZyme expression varied depending on the carbon substrate available to this fungal isolate. DESeq2 (Love et al. 2014) analysis showed that the highest levels of differentially expressed CAZymes are found when comparing reed canary grass (the most recalcitrant substrate tested) to the other less complex carbon sources (filter paper, cellobiose, and glucose) (Figure 5, Supplemental Tables 3–14). Filter paper is more difficult to break down than cellobiose or glucose, which was reflected in hundreds of differentially expressed predicted CAZymes when comparing filter paper to cellobiose or glucose. Cellobiose, with the closest structural similarity to glucose of the substrates, showed the most similar pattern of CAZyme expression to glucose.

**Figure 5.** Numerous predicted CAZymes are differentially regulated across different carbon sources in *N. cameroonii* var. constans. For each substrate comparison, the number of genes containing at least one of the six CAZyme domains is enumerated. Genes are only included if they are significantly differentially expressed (padj<0.05) and have an average TPM greater than 2 for one or both conditions compared.

Using DESeq2 (Love et al. 2014), we identified significantly differentially expressed genes for the different comparative growth conditions. Gene expression of *N. cameroonii* var. constans grown on reed canary grass, filter paper, and cellobiose was compared to that on glucose as a baseline. The number of significantly upregulated genes was highest in the reed canary grass/glucose comparison, followed by filter paper/glucose and then cellobiose/glucose. This is observed in both the all-genes dataset (Figure 6, Supplemental Figures 3–5) and the CAZymes-only dataset (Supplemental Figure 7).

**Figure 6.** Differential expression analysis of *N. cameroonii* var constans grown on cellobiose, filter paper (FP), or reed canary grass (RCG), each compared to glucose. The top 100 predicted genes based on the lowest p-values across any of the comparisons are shown in this heatmap. Colors correspond to the log2 fold change value of a predicted gene for each condition. For the genes annotated as CAZymes, domains are listed, with their predicted counts within that gene. For example, “Dockerin x3” indicates that 3 dockerin domains were identified by dbCAN3 in the associated gene. Gene ordering is based on expression similarity. ProteinID refers to the annotated MycoCosm designation.

When comparing each growth substrate, pairwise comparisons showed substantially more differentially expressed genes between reed canary grass and each of the other substrates than pairwise comparisons among filter paper, cellobiose, and glucose (Supplemental Figures 8 and 9). Approximately 53–64% of expressed genes (with an average TPM>2 for at least one condition) were significantly differentially expressed between reed canary grass and each of the other substrates, while only 8–28% were significantly differentially expressed for pairwise comparisons between filter paper, cellobiose, and glucose (Supplemental Figure 9). In pairwise comparisons with reed canary grass, the majority (∼60–65%) of significantly differentially expressed genes were upregulated in reed canary grass cultures (Supplemental Figure 9). These results confirm that more genes are upregulated in breaking down a complex lignocellulosic material and utilizing its components. However, it is important to note that reed canary grass cultures also contained the highest number of significantly *downregulated* genes relative to the other substrates (Supplemental Figure 9) and interestingly, these downregulated genes also include numerous CAZymes (Supplemental Table 15). These observations are consistent with large-scale changes in gene regulation patterns occurring when grown on complex lignocellulosic material.

### *N. cameroonii* var. constans and Methanogen Co-cultures Convert Lignocellulose to Methane

*N. cameroonii* var. constans was isolated from a previously characterized microbial community, where the community produced methane as a major end product and maintained its membership and metabolic output for multiple years of cultivation (Peng et al. 2021). A methanogen from this microbial community (*Methanobrevibacter* sp.) was separately isolated as well (Supplemental Table 16). Anaerobic fungi and methanogens exist in commensal and symbiotic relationships in the rumen, with hydrogen produced by fungi being converted to methane by methanogenic archaea such as in natural consortia (Jin et al. 2011; Gilmore et al. 2019) and synthetic co-cultures (Swift et al. 2019; Leggieri et al. 2021). Owing to their high abundances within their respective phyla in these goat fecal communities, we hypothesized that *N. cameroonii* var. constans and the lab-isolated methanogen, *Methanobrevibacter* sp., are especially capable of synergy relative to other anaerobic fungi – methanogen pairs. Methane production was measured for fungal-methanogen co-cultures: 4 different methanogens were paired with *N. cameroonii* var. constans and *N. californiae* (Figure 7). Productive partnership (evidenced by methane detection) was observed for all tested samples via the generation of methane from lignocellulosic substrates. Differences in measured gas were variable and can be ascribed to stochasticity in relative growth rates between strains of anaerobic gut fungi and methanogens; nevertheless, we observed similar amounts of produced methane at late co-culture times. These results are consistent with those observed by Peng *et al*. 2021, in which membership by both methanogenic archaea and fungi were important for methane production. *N. cameroonii* var. constans is therefore a versatile chassis organism– the strain can be cultivated in communities, both natural and synthetically designed to capture metabolic interactions found in the rumen.

**Figure 7.** Methane production of methanogens when co-cultured with *N. californiae* and *N. cameroonii* var. constans shows ability of anaerobic fungi to degrade lignocellulose into fermentation products that methanogens can convert to methane. “*M. sp*.” is the *Methanobrevibacter sp.* isolated from the same natural microbial community as *N. cameroonii* var. constans. Note that methane was not detected (N.D.) fungal monocultures. Error bars represent standard deviation of three biological replicates.

## Conclusions

This new isolate, *N. cameroonii* var. constans, is a robust anaerobic fungal strain with a considerable CAZyme portfolio, with potentially many more enzymes involved in degradation than previously characterized. Structure-based prediction with AlphaFold revealed diverse, unannotated, catalytic domains motivating future experimental validation. This strain’s ease of cultivation is supported by its growth on a variety of different substrates, including reed canary grass, filter paper, cellobiose, and glucose (Supplemental Figure 10), with transcriptional analyses showing a clear impact of substrate on gene expression. Additionally, its ability to exist long-term in stable, naturally-derived consortia as well as pair with methanogens in synthetic communities helps elucidate its potential for further microbial community studies, to reveal complex metabolic and spatial interactions between anaerobic fungi and other rumen members. This novel strain is a strong candidate for an anaerobic fungal chassis, and as such, it offers a step towards illuminating complicated anaerobic microbial interactions and substrate degradation mechanisms.

## Data Availability

The genome assembly and annotations are available from MycoCosm (Grigoriev et al. 2014) (https://mycocosm.jgi.doe.gov/Neocon1) and have been deposited at DDBJ/ENA/Genbank under the accession (TO BE PROVIDED UPON PUBLICATION). The transcriptome data have been deposited to SRA under sequencing project IDs 1304688 and 1304689. Additional supporting files for RNAseq analyses and CAZyme annotations are available at https://github.com/O-Malley-Lab/N_var_constans.

## Acknowledgements

We acknowledge funding support from the Department of Energy, Office of Science (DE-SC0020420 and DE-SC0022142), the Institute for Collaborative Biotechnologies (W911NF-19-D-0001 and W911NF-19-2-0026), and National Science Foundation (2128271). This work was also funded by the of the Department of Energy (DOE) Joint BioEnergy Institute (http://www.jbei.org) supported by the Office of Biological and Environmental Research of the DOE Office of Science through contract DE-AC02–05CH11231 between Lawrence Berkeley National Laboratory. The work (proposal: 10.46936/10.25585/60001061) conducted by the U.S. Department of Energy Joint Genome Institute (https://ror.org/04xm1d337), a DOE Office of Science User Facility, is supported by the Office of Science of the U.S. Department of Energy operated under Contract No. DE-AC02-05CH11231.

The authors would like to thank Asaf Salamov for useful discussions regarding bioinformatics, Bernard Henrissat and CAZy team for CAZyme annotations, and Frank Kinnaman and David Valentine for gas chromatography assistance and helpful discussions. The authors are grateful for support from the Joint Genome Institute and Arizona Genome Institute in this work. The work (proposal: 10.46936/10.25585/60001061) conducted by the U.S. Department of Energy Joint Genome Institute (https://ror.org/04xm1d337), a DOE Office of Science User Facility, is supported by the Office of Science of the U.S. Department of Energy operated under Contract No. DE-AC02-05CH11231. We acknowledge Francis Martin and JGI for the genome sequence for *M. rebaudengoi* and Kevin Solomon and JGI for the genome sequence of *Piromyces sp. UH3-1*.

Use was made of computational facilities purchased with funds from the National Science Foundation (CNS-1725797) and administered by the Center for Scientific Computing (CSC). The CSC is supported by the California NanoSystems Institute and the Materials Research Science and Engineering Center (MRSEC; NSF DMR 2308708) at UC Santa Barbara. The authors also acknowledge the use of the Biological Nanostructures Laboratory within the California NanoSystems Institute, supported by the University of California, Santa Barbara and the University of California, Office of the President.

## Conflicts of interest

The authors declare no conflicts of interest.

